# Transient Suppression of Dopamine Transporter Palmitoylation by Methamphetamine: Implications for Transport Regulation

**DOI:** 10.1101/2025.05.02.651918

**Authors:** Moriah J. Hovde, Danielle E. Bolland, Corey D. Kleinsasser, Madhur Shetty, Aaron C. Blackwell, Mikhail Y. Golovko, Svetlana A. Golovko, Christopher R. Brown, James D. Foster, Roxanne A. Vaughan

**Author notes:** To whom correspondence should be addressed: Roxanne A. Vaughan, Ph.D. Department of Biomedical Sciences, University of North Dakota School of Medicine and Health Sciences, Grand Forks, ND 58202-9037., James D. Foster, Ph.D., Department of Biomedical Sciences, University of North Dakota School of Medicine and Health Sciences, Grand Forks, ND 58202-9037. MJ Hovde: Massachusetts General Hospital Institute for Neurodegenerative Disease, Harvard Medical School, Charlestown, MA 02129; DE Bolland: The University of Minnesota Morris, Division of Science and Math, Biology Discipline. Morris, MN 56267.

## Abstract

The dopamine transporter (DAT) exerts temporal and spatial control over dopaminergic neurotransmission through reuptake of extracellular dopamine (DA). The functional capacity of DAT is under the control of signaling inputs and post-translational modifications that confer acute presynaptic regulation of reuptake in response to physiological needs, and dysregulation of these processes may contribute to DA imbalances in mood disorders and drug addiction. A key modification of DAT is palmitoylation, a lipid adduction that enhances transport velocity, is suppressed by protein kinase C, and opposes protein kinase C-mediated down-regulation. Here we now show in rat striatum and heterologous cells that transporter palmitoylation is also linked to methamphetamine (METH), undergoing rapid and transient reductions in response to the drug. The time course and other characteristics of palmitoylation reduction parallel those of METH-induced transport down-regulation, and a palmitoylation-deficient DAT mutant shows enhanced down-regulation to METH, supporting a mechanistic link between reduction of the modification and reduced reuptake activity. Recovery rates differed, however, with palmitoylation returning to starting levels more rapidly than reuptake, indicating that down-regulation mechanisms remain engaged with transporters that have undergone repalmitoylation. These results support palmitoylation as a rapid response mechanism that modulates DAT entry into METH-induced down-regulation states and suggest a broader role for the modification in control of reuptake in additional physiological and pathophysiological conditions.

## INTRODUCTION

The dopamine transporter (DAT) is expressed on plasma membranes of axons and terminals of midbrain dopaminergic neurons that project to the basal ganglia and cortical areas of the brain (Nirenberg, Vaughan et al. 1996). DAT plays a key role in dopamine (DA) neurotransmission in these regions through reuptake of extracellular transmitter, and dysregulation of DAT function may result in dopaminergic imbalances in mood, psychiatric, and movement disorders (Pramod, Foster et al. 2013). Many abused drugs including transport blockers such as cocaine and transported substrates such as amphetamine (AMPH) and methamphetamine (METH) act at DAT to inhibit DA reuptake, elevating transmitter to supraphysiological levels that underlie psychomotor stimulation and addiction (Chen and Reith 2000, Gowrishankar, Hahn et al. 2014).

In addition to their pharmacological actions, AMPH and METH affect DAT physiologically by inducing kinetic and endocytotic down-regulation processes that persist after drug has been cleared, which further increases the magnitude and duration of transport reductions and contributes to addictive potential (Schmitt and Reith 2010, Vaughan and Foster 2013, Bu, Farrer et al. 2021). Under normal conditions DAT activity and surface expression are regulated by multiple signaling systems that synergistically function to coordinate transmitter clearance with physiological needs (Vaughan and Foster 2013). A major pathway linked to transport reduction and endocytosis is protein kinase C (PKC), and the similarities of AMPH-, METH-, and PKC-induced down-regulation events have drawn attention to post-translational modifications in psychostimulant mechanisms. DAT is a target of PKC, which stimulates phosphorylation of N-terminal domain residues that mediate reduced reuptake (Huff, Vaughan et al. 1997, Vaughan, Huff et al. 1997, Granas, Ferrer et al. 2003, Cervinski, Foster et al. 2005, Foster, Yang et al. 2012). Phosphorylation of this domain is also stimulated by AMPH and METH in a PKC-dependent manner, implicating the kinase as a key element in psychostimulant substrate actions (Huff, Vaughan et al. 1997, Vaughan, Huff et al. 1997, Granas, Ferrer et al. 2003, Cervinski, Foster et al. 2005, Foster, Yang et al. 2012).

A related modification of DAT controlled by PKC is palmitoylation, the reversible adduction of a lipid moiety to sulfhydryl groups of cytoplasmically-oriented cysteines. In rat (r) and human (h) transporters palmitate incorporation occurs on multiple sites, including Cys580/581 at the interface between the intracellular end of transmembrane domain 12 (TM12) and the cytoplasmic C-terminus (Foster and Vaughan 2011). Palmitoylation increases DA transport velocity, opposes PKC-induced down-regulation, and is suppressed by activation of PKC (Foster and Vaughan 2011, Moritz, Rastedt et al. 2015, Bolland, Moritz et al. 2019). These characteristics are opposite those of transporter phosphorylation and the modifications thus function synergistically to integrate incoming information in control of reuptake.

In this study we now show that DAT palmitoylation is also mechanistically linked to METH. For these studies, we utilized a single high-dose injection model in rats that has been extensively characterized by Fleckenstein and colleagues to investigate mechanisms of METH-induced transport down-regulation (Fleckenstein, Metzger et al. 1997, Kokoshka, Vaughan et al. 1998). These studies showed that METH, but not cocaine, induces transport reductions that are rapid, reversible, and not due to changes in DAT protein levels, consistent with a post-translational mechanism. Related *in vitro* studies demonstrating transport down-regulation to METH in cultured cells or synaptosomes support that responses do not require neuronal circuitry or input from DA receptors and are PKC-dependent (Fleckenstein, Metzger et al. 1997, Kokoshka, Vaughan et al. 1998, Metzger, Haughey et al. 2000, Sandoval, Hanson et al. 2000, Sandoval, Riddle et al. 2001, Cervinski, Foster et al. 2005, German, Hanson et al. 2012).

Using this injection paradigm, we now show that palmitoylation of striatal transporters undergoes rapid and transient reductions in response to METH, which to the best of our knowledge is the first demonstration of *in vivo* reversibility of a DAT post-translational modification following an acute stimulus. Palmitoylation reductions occurred within minutes of METH injection, similar to the time course of transport down-regulation, and in model cells palmitoylation-deficient C580A DAT displayed a greater magnitude of METH-induced transport down-regulation than WT DAT. These findings, in conjunction with other similarities including PKC dependency and lack of cocaine effect, support that palmitoylation functions to oppose entry of DAT into METH-induced down-regulated states. Recovery of transport was markedly slower than repalmitoylation, however, potentially indicating that palmitoylation status exerts minimal effect on reversal of down-regulation processes. These findings support palmitoylation as a rapid response mechanism that modulates DAT outcomes to METH and suggest palmitoylation more broadly as a regulator of reuptake in additional physiological and pathophysiological conditions.

## MATERIALS AND METHODS

### Materials

[7,8-^3^H]DA (45 Ci/mmol) was from ViTrax (Placentia, CA, USA); DAT monoclonal antibody 16 (MAb 16) was previously authenticated (Gaffaney and Vaughan 2004) and available commercially (Invitrogen-ThermoFisher MA5-24796); N-ethylmaleimide (NEM), high capacity NeutrAvidin-agarose resin, and bicinchoninic acid (BCA) protein assay reagent were from ThermoFisher Scientific (Waltham, MA, USA); HPDP-biotin was from APExBIO (Houston, TX, USA); (−)-Cocaine, AMPH, METH, DA, and other fine chemicals were from MilliporeSigma (Sheboygan Falls, WI, USA). Bisindolylmaleimide I (BIM I) was from Cayman Chemical (Ann Arbor, MI, USA). Rats were purchased from Envigo (Lafayette, IN, USA). All animals were housed and treated in accordance with regulations established by the National Institutes of Health and approved by the University of North Dakota Institutional Animal Care and Use Committee.

### Drug Injection and Membrane/Synaptosome Preparation

Male Sprague-Dawley rats (175-300 g) were given single 1 ml subcutaneous injections of saline, METH, or (−)-cocaine to achieve final drug dosages of 15 mg/kg. Animals were decapitated at indicated times after injection and striata were rapidly removed, weighed, and placed in ice-cold sucrose phosphate buffer (SP, 0.32M sucrose, 10 mM Na_2_PO_4_, pH 7.4). To prepare synaptosomes tissue was homogenized at 4 °C in 2 ml of SP buffer using a glass/Teflon homogenizing apparatus, volume brought up to 10 ml, and homogenates centrifuged at 3,000 x g for 3 min at 4 °C to remove nuclei and debris. The supernatant fraction was transferred to a fresh tube and centrifuged at 17,000 x g for 12 min. The pellet was washed two times by resuspension in 5 ml of SP buffer and centrifugation at 17,000 x g for 12 min at 4 °C. The final pellet was resuspended with SP buffer at 15 mg/ml original striatal wet weight and synaptosomes analyzed for [^3^H]DA uptake or DAT immunoblotting. For analysis of palmitoylation, striatal membranes were prepared by homogenization of striata in SP buffer containing 100 mM EDTA (SP+) using a Polytron tissue homogenizer and centrifugation at 1,000 x g for 10 min at 4 °C to pellet nuclei and debris. The supernatant fraction was transferred to a fresh tube and centrifuged at 12,000 x g for 12 min and the pellet was resuspended in 5 ml of SP+ buffer and centrifuged at 12,000 x g for 12 min at 4 °C. The final membrane pellet was resuspended with SP+ buffer at a final concentration of 50 mg/ml original striatal wet weight and used for palmitoylation analysis via ABE.

### Cell Culture and Treatments

Lilly laboratory porcine kidney (LLC-PK_1_) cells expressing rDAT were grown in alpha minimum essential medium (AMEM) supplemented with 5% fetal bovine serum, 2 mM L-glutamine, 200 μg/ml G418, 100 μg/ml penicillin/streptomycin, and 0.25 µg/ml amphotericin B and maintained in a humidified 5% CO_2_ environment at 37 °C. For analysis of METH, AMPH, cocaine, or BIM I effects, cells were treated for the times or combinations described in the text and figure legends with vehicle, 10 μM METH, 10 µM AMPH, 10 μM (−)-cocaine, or 10 μM BIM I. For BIM I treatments, cells were preincubated with BIM I for 5 min prior to the addition of METH and continued incubation for 30 additional min. At the end of the treatment time, the medium was removed, cells were washed with the indicated buffer and analyzed for DAT palmitoylation or [^3^H]DA uptake as described below.

### [^3^H]DA Uptake Analysis

Uptake in synaptosomes was initiated by the addition of 100 μl of synaptosomes to 900 μl of modified Krebs phosphate buffer (MKP, 16 mM potassium phosphate, 126 mM NaCl, 4.8 mM KCl, 1.4 mM MgSO_4_, 10 mM glucose, 1.1 mM ascorbic acid, and 1.3 mM CaCl_2_, pH 7.4) bringing the final concentration of [^3^H]DA to 10 nM and total DA to 1 µM. For DA saturation analyses, total DA concentrations were between 0.3 to 1 µM containing 10 nM [^3^H]DA. Assay tubes were incubated for 5 min with shaking at 30 °C, and transport was stopped by the addition of 5 ml ice-cold SP buffer and filtration through a glass/fiber filter using a Brandel filtration device. Radioactivity remaining on washed filters was measured by liquid scintillation counting. Nonspecific uptake was determined by the addition of 100 μM (−)-cocaine and subtracted from total uptake to determine specific uptake values. K_m_ and V_max_ constants were determined by nonlinear regression analysis using GraphPad Prism Software. For analyses in cells WT or mutant rDAT LLC-PK_1_ cells were grown in 24-well plates to 80% confluence and rinsed twice with 0.5 ml of 37 °C KRH buffer. Cells were then incubated with vehicle, 10 µM METH, or 10 µM (−)-cocaine in Krebs-Ringer/HEPES buffer (KRH: 25 mM HEPES, 125 mM NaCl, 4.8 mM KCl, 1.2 mM KH_2_PO_4_, 1.3 mM CaCl_2_, 1.2 mM MgSO_4_, 5.6 mM glucose, pH 7.4) buffer for the indicated times. For washout studies after METH treatment, cells were rapidly washed with 37 °C KRH to remove the drug followed by incubation with KRH for the indicated times at 37 °C. DA uptake assays were initiated by the addition of 10 µl of a 50X DA stock solution to bring the final concentration of [^3^H]DA to 10 nM and total DA to 3 µM. Uptake was conducted for 8 min and terminated by rapidly washing the cells two times with ice-cold KRH buffer. Nonspecific uptake was determined by the addition of 100 μM (−)-cocaine to parallel wells and subtracted from total uptake values to determine specific uptake. Cells were lysed with 1% Triton X-100 and radioactivity in the lysates was measured by liquid scintillation counting.

### Cell Membrane Preparation

Cells were grown to approximately 90% confluency in 15 cm cell culture dishes, treated with the indicated drug or vehicle for the indicated time, media was removed, and cells were washed twice with ice-cold Buffer B (0.25 M sucrose, 10 mM triethanolamine, pH adjusted to 7.8 with 100 mM acetic acid, and supplemented with 1 μM phenylmethylsulphonyl (PMSF) and 5 μM ethylenediaminetetraacetic acid (EDTA). For washout studies, cells were rapidly washed with 37 °C KRH buffer to remove the drug. After the initial rapid wash, the cells were incubated with KRH for the indicated times at 37°C. The cells were then placed on ice and washed twice with ice-cold Buffer B. The cells were then scraped using Buffer B, collected in 1.7 ml microcentrifuge tubes, and centrifuged at 3,000 x g for 5 min at 4 °C. The supernatant was removed and the cell pellet gently resuspended in Buffer C (0.25 M sucrose, 10 mM triethanolamine, 1 mM EDTA, pH 7.8 with 100 mM acetic acid, supplemented with 1 μM PMSF and 5 μM EDTA), transferred to a Dounce homogenizer and homogenized with 30 up and down strokes. The sample was then centrifuged for 10 min at 800 x g to remove nuclei and cell debris. The supernatant was then collected and centrifuged at 16,000 x g for 12 min, and the resulting membrane pellet was resuspended in 1 ml SP buffer (0.32 M sucrose, 10 mM sodium phosphate, pH 7.4 with 1 μM PMSF and 5 μM EDTA), assayed for protein concentration and stored at −20 °C for analysis.

### Acyl-Biotin Exchange

Palmitoylated proteins were detected using procedures adapted from Wan et al. (Wan, Roth et al. 2007). (i) To block free cysteine thiol groups on DAT, cell membranes (200-300 µg protein) were solubilized in 250 µl lysis buffer (50 mM HEPES pH 7.0, 2% SDS (w/v), 1 mM EDTA) containing 25 mM NEM and incubated for 20 min in a 37 °C water bath followed by end-over-end mixing for at least 1 h at ambient temperature. Proteins were precipitated by the addition of 1 mL acetone and mixing, followed by centrifugation at 18,000 x g for 10 min. The protein pellet was resuspended in 250 µl lysis buffer containing 25 mM NEM and incubated for an hour at ambient temperature with end-over-end mixing. This process was repeated a final time with overnight incubation and end-over-end mixing. NEM was removed by acetone precipitation and centrifugation, and the protein pellet was resuspended in 250 μL 4SB buffer (50 mM Tris, 5 mM EDTA, 4% SDS, pH 7.4). Final protein pellets were resuspended in 200 μL 4SB buffer. (ii) Endogenous thioester-linked palmitoyl groups on DAT were removed by treatment with hydroxylamine (HA), using treatment of parallel aliquots with Tris-HCl buffer as a negative control to verify NEM blockade of free sulfhydryl groups. The resuspended protein sample is split into equal volumes (100 µl) that are treated with 50 mM Tris-HCl, pH 7.4 or 0.7 mM HA in Tris buffer adjusted to pH 7.4, and incubated at room temperature for 30 min with end-over-end mixing. (iii) Both samples are then treated with sulfhydryl-specific HPDP-Biotin (0.4 mM final, 50 mM Tris-HCl, pH 7.4) for 1 h at room temperature with end-over-end mixing to biotinylate free sulfhydryl groups liberated by HA treatment. HA and unbound biotin were removed with three sequential acetone precipitations (centrifugation at 18,000 x g for 10 min, supernatant fraction removal, and resuspension of the protein pellet in 250 µl 4SB) followed by resuspension/solubilization of the final protein pellet in 75 µl of lysis buffer. (iv) For determination of total DAT content in the ABE sample, a 10 μL aliquot was set aside for immunoblotting while 65 μL was diluted in 1500 μl Tris buffer and incubated with 50 μL of a 50% slurry of high-capacity NeutrAvidin resin to affinity purify the biotinylated proteins overnight at 4°C with end-over-end mixing. Unbound proteins were removed by three cycles of 8,000 x g centrifugation, removal of the supernatant fraction, and resuspension in 750 μL radioimmunoprecipitation assay buffer (RIPA: 1% Tx-100, 1% sodium deoxycholate, 0.1% SDS, 125 mM sodium phosphate, 150 mM NaCl, 2 mM EDTA, 50 mM NaF). Proteins were eluted from the final pellet by incubation in 2x Laemmli sample buffer (SB: 125 mM Tris-HCl, 20% glycerol, 4% SDS, 200 mM DTT, 0.005% bromophenol blue) for 20 min at ambient temperature. Eluted samples were then subjected to SDS-PAGE and immunoblotted for rDAT with mouse MAb16 primary antibody.

### Quantification of DAT Palmitoylation by ABE

Palmitoylation of DAT was quantified as previously described (Moritz, Rastedt et al. 2015, Brown and Foster 2023, Brown, Shetty et al. 2025). Briefly, aliquots (10 µl) taken from final resuspension (75 µl) just prior to NeutrAvidin extraction in the ABE assay were used to directly assess total DAT levels in the sample and normalize DAT palmitoylation level in this same sample for comparison to other samples and treatments. Palmitoylated DAT band intensities were quantified using Quantity One software (Bio-Rad), normalized to total DAT protein present, and expressed as % control. Bands in immunoblots represent 13.3% of the total and 86.7 % of the palmitoylated DAT in each sample. Control values were set to 100%.

### Immunoblotting

Samples were resolved using 4-20% SDS-polyacrylamide gels, transferred to PVDF membranes and assayed for DAT as previously described (Gaffaney and Vaughan 2004). Membranes were probed using a 1:1000 dilution of mouse monoclonal N-terminal Ab16 (MAb16) for rDAT (ThermoFisher Scientific). Anti-mouse alkaline phosphatase-conjugated secondary antibody was utilized along with ImmunStar (Bio-Rad) substrate and the Bio-Rad gel documentation system to visualize the blots and quantify levels of immunostaining in the linear range using Quantity One software (Bio-Rad).

### METH Extraction and LC-MS/MS Analysis

Synaptosome samples (25 µl, n=6) taken 30 min after METH injection were spiked with a deuterium-labeled METH as an internal standard (100 pg methamphetamine-d_5_) followed by methanol extraction by addition of 75 µl methanol, thorough mixing and incubation in a sonic water bath for 1 min, followed by centrifugation at 20,000 x g for 10 min at room temperature. The supernatant fraction was transferred to a sample vial. Five µL of sample was loaded onto an ACQUITY UPLC HSS T3 column (1.8 μM, 100 Å pore diameter, 2.1 × 150 mm) with an ACQUITY UPLC HSS T3 precolumn (1.8 μM, 100 Å pore diameter, 2.1 × 5 mm, Waters, Milford, MA) and METH was resolved using a Waters Acquity UPLC system (Waters, Milford, MA). A gradient elution with solvent A (0.1% formic acid in water) and solvent B (0.1% formic acid in acetonitrile) at 0.3 mL/min was used. Initial 10% B was maintained for 0.5 min, then increased to 40% over 3.5 min. At 6.5 min, %B was increased to 75%, and at 9 min returned back to 10%. The column was equilibrated with 10% B for 3 min between injections. MS/MS analysis was performed on a Waters Xevo TQ-S triple quadrupole mass spectrometer (Waters, Milford, MA) using multiple reaction monitoring mode. MS was operated in a positive ESI mode. METH was quantified against the deuterium-labeled internal standard METH-d_5_ based on the calibration curve and the mean values used to calculate the residual METH in the assay tube. The following mass transitions (with collision energies indicated in parentheses, V) were used: for METH-d_5_ - 155.18/120.96 (10, for confirmation) and 155.18/91.73 (16, for quantification); for METH – 150.15/118.97 (10, for confirmation) and 150.15/90.96 385.37/95.10 (16, for quantification). MassLynx 4.1 (Waters) was used for instrument control and data processing.

## RESULTS

To determine the effects of *in vivo* METH or (−)-cocaine on DAT palmitoylation male Sprague-Dawley rats were given single s.c. injections of saline, 15 mg/kg METH or 15 mg/kg (−)-cocaine, sacrificed between 10-60 min post-injection, and striatal transporters analyzed for palmitoylation using the acyl biotinyl exchange (ABE) assay (**Figure 1**). In this method endogenous palmitate groups are chemically replaced with a biotinylated moiety that allows for extraction and quantitative immunoblotting of originally palmitoylated forms, permitting assessment of real-time *in vivo* changes in modification levels. **Figure 1A** shows that within 10 min of METH injection, DAT palmitoylation was reduced to 66.8 ± 10.1% of control levels (p<0.05) and remained suppressed through 30 min (50.8 ± 0.7% of control, p<0.001) and 60 min (48.6 ± 0.7% of control, p<0.001). In separate experiments we investigated uptake blocker actions using (−)-cocaine, and found that 30 min after injection, DAT palmitoylation was unchanged (94.8 ± 5.6% of control, p>0.05) compared to that of rats given METH in parallel that showed reductions to 76.6 ± 5.5% of control (p<0.05) (**Figure 1B**). Equivalent levels of total DAT were present in all ABE samples, indicating that palmitoylation reductions are due to enzymatic palmitate alterations and not to changes in DAT expression.

**Figure 1.**
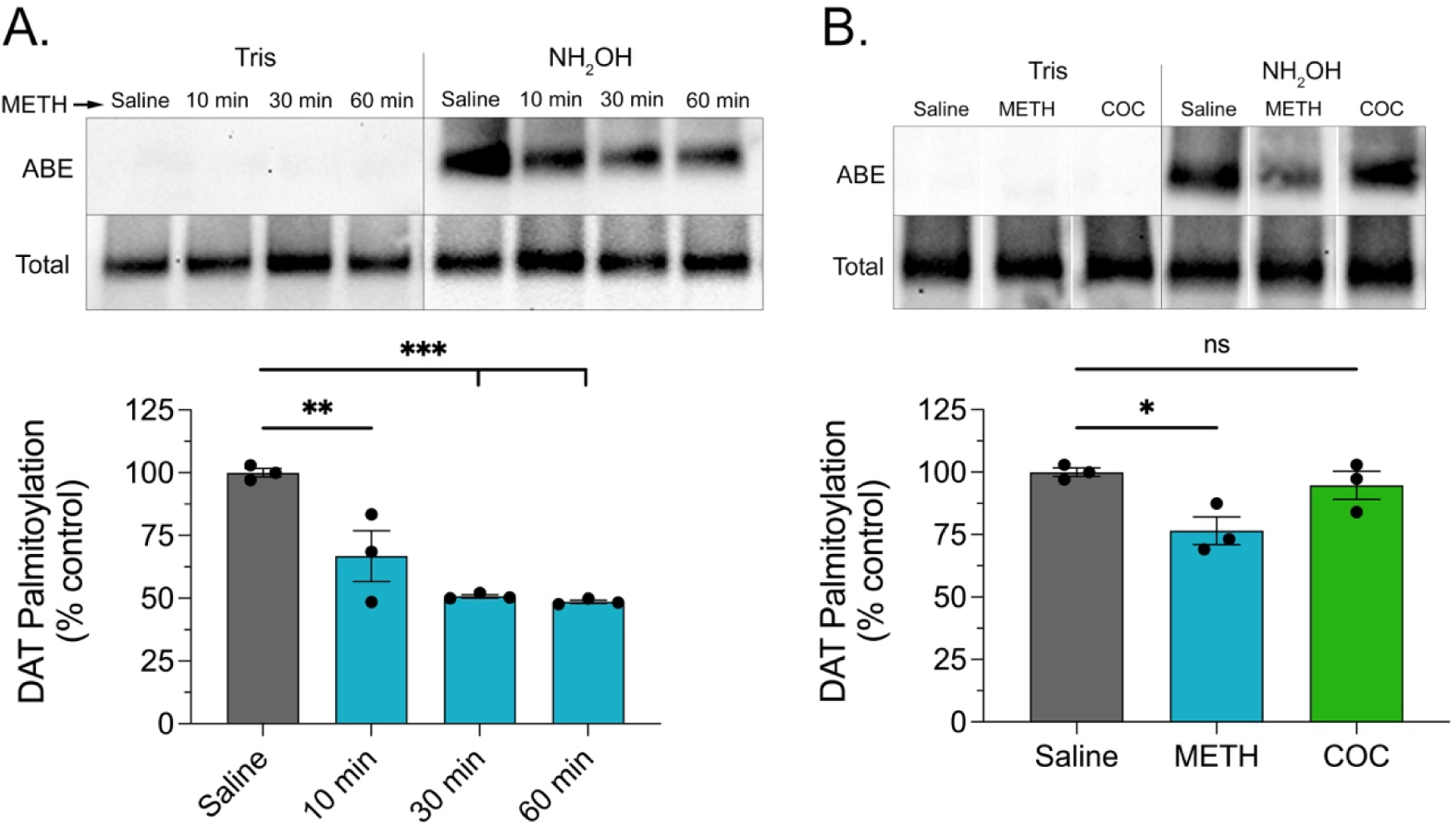
DAT palmitoylation is reduced by *in vivo* METH. Male Sprague-Dawley rats were given s.c. injections of saline, METH (15 mg/kg), or (−)-cocaine (15 mg/kg) and sacrificed at indicated times (A) or after 30 min (B). Striatal tissue was isolated and analyzed by ABE for DAT palmitoylation (upper rows) or by immunoblot for total DAT (lower rows). Blots show ABE, Tris specificity controls, and total DAT analyses from representative experiments, and histograms show quantification of DAT palmitoylation normalized for total DAT protein and expressed as a fraction of saline control set to 100% (means ± S.E. of 3 independent experiments performed in duplicate). * p<0.05; **, p<0.01; ***, p<0.001 vs. control; one-way ANOVA with Tukey’s post hoc test. n.s., not significant. White spaces between lanes in panel B indicate cropping to remove duplicate samples.

To further characterize the response, we analyzed the additional time points shown in **Figure 2**. Here rats were given s.c. injections of saline or 15 mg/kg METH and sacrificed between 5-150 min post-injection. Within 5 min DAT palmitoylation was reduced to 75.5 ± 3.2% of control, with decreases maintained through 10 min (68.7 ± 7.4% of control), 30 min (59.0 ± 4.1% of control), and 60 min (63.3 ± 6.7% of control) (all p<0.001 or p<0.0001). At later time points palmitoylation gradually returned to starting levels, reaching values of 86.4 ± 2.8% of control at 90 min, 103.2 ± 2.3% of control at 120 min, and 100.7 ± 4.8% of control at 150 min (all p>0.05 vs. control and p<0.001-0.0001 vs. palmitoylation level at 30 min). The rapid reduction of palmitoylation by METH closely correlates with drug entry into the brain, which occurs within minutes (Gorman, Coward et al. 2023), and the return of palmitoylation to control levels could potentially follow from metabolic clearance of the drug, which in rats occurs with a T_½_ of ∼1 h (Cho, Melega et al. 2001, Segal and Kuczenski 2006). In addition, the lack of effect from cocaine supports that the reductions do not follow solely from drug-induced elevations in extracellular DA.

**Figure 2.**
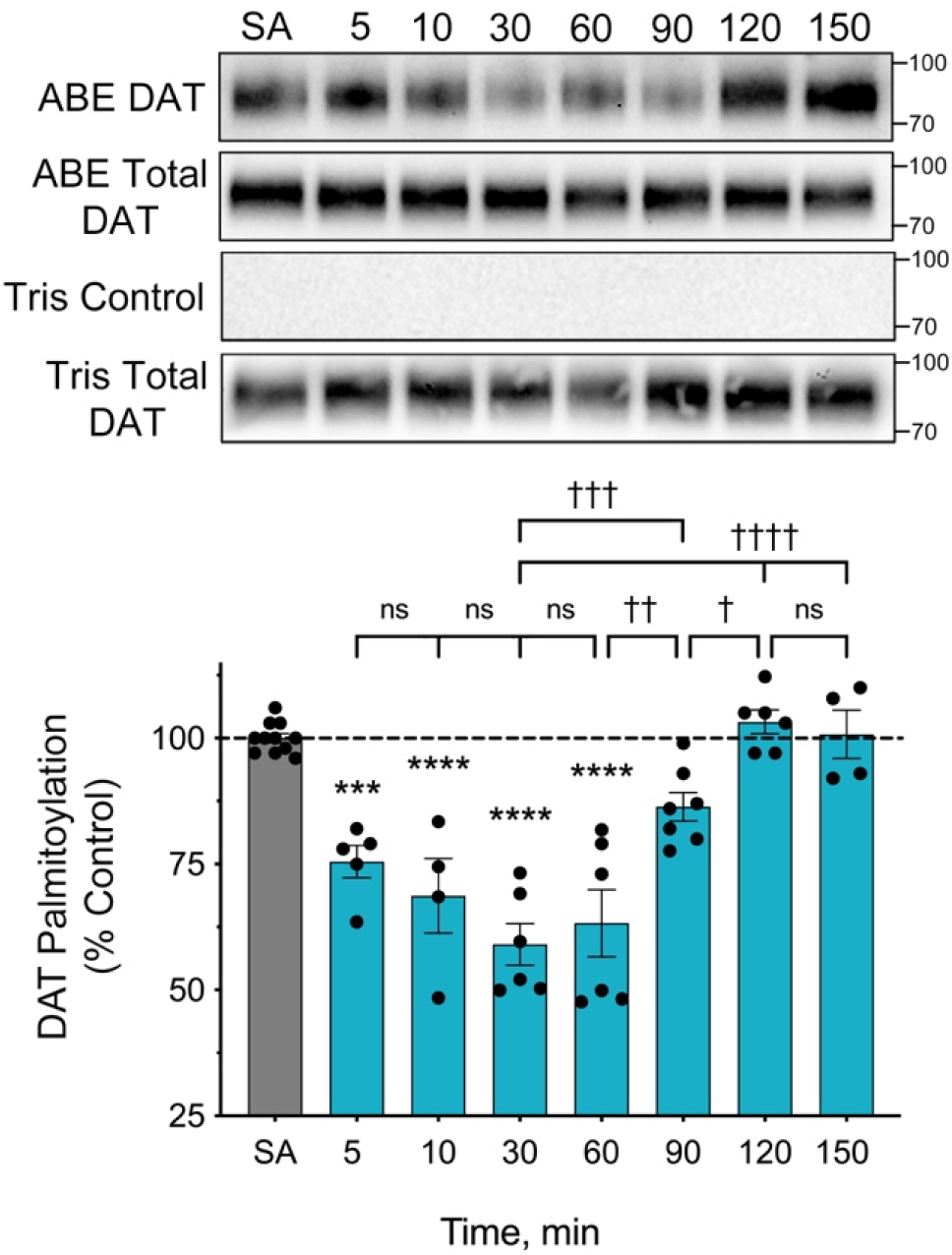
Onset and reversibility of DAT palmitoylation reductions to *in vivo* METH. Male Sprague-Dawley rats were given s.c. injections of saline (SA) or METH (15 mg/kg), sacrificed at the indicated times, and striatal membranes analyzed for DAT palmitoylation. The blots show ABE (top row), corresponding total DAT (second row), and ABE specificity controls and totals (bottom two rows) from a representative experiment. Histogram shows quantification of DAT palmitoylation normalized for total DAT expressed as a fraction of saline control set to 100% (means ± S.E. of 4-7 experiments). ***, p<0.001; ****, p<0.0001; indicated time points vs. SA control; ^†^, p<0.05; ^††^, p<0.01; ^†††^ p<0.001, ^††††^, p<0.0001, confidence values between indicated time points. n.s, not significant, (one-way ANOVA with Tukey’s post hoc test).

We then examined the response of transport activity to METH treatment across the time points in which palmitoylation showed suppression and recovery (**Figure 3**). For these studies rats were given s.c. injections of saline or 15 mg/kg METH and sacrificed 10-150 min after injection. Synaptosomes were prepared from striatal tissue and washed two times during preparation, which was previously demonstrated to reduce residual METH to levels that do not pharmacologically inhibit uptake (Sandoval, Hanson et al. 2000). We also used LC-MS/MS to quantify METH levels and in six independent samples collected 30 min after injection confirmed amounts to be consistent with assay concentrations of 0.4-3 nM, below the pharmacological threshold for uptake inhibition.

**Figure 3.**
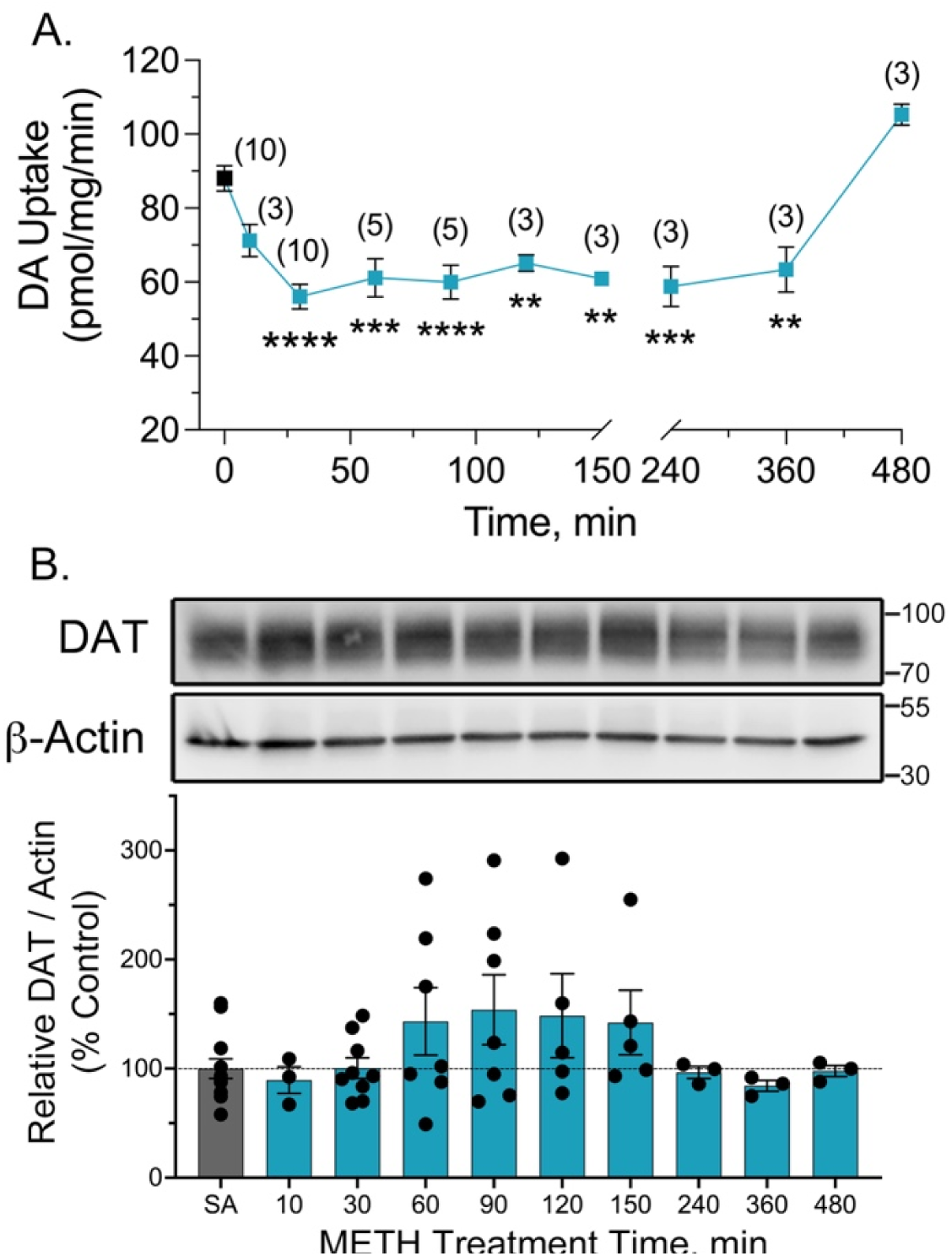
Onset and reversibility of DA transport reductions in synaptosomes from METH-treated rats. Male Sprague-Dawley rats were given s.c. injections of saline or METH (15 mg/kg) and sacrificed at the indicated time points. Synaptosomes were prepared from striatal tissue collected at each time point and equal amounts assessed for [^3^H]DA uptake or immunoblotted for total DAT. (A). DA transport at indicated times after injection (means ± S.E. of 3-10 experiments performed in quadruplicate; number of independent experiments indicated in parentheses). **, p<0.01, *** p<0.001, ****, p<0.0001 relative to control (one-way ANOVA with Tukey’s post hoc test). (B). Representative immunoblots of total DAT and β-actin from aliquots of synaptosomes utilized for uptake. Histogram shows quantification of DAT normalized to β-actin for each time point (means ± S.E. of 4-7 experiments). All values p>0.05 vs control. (one-way ANOVA with Tukey’s post hoc test)

As previously described (Fleckenstein, Metzger et al. 1997, Kokoshka, Vaughan et al. 1998), METH injection induced a rapid decrease in synaptosomal [^3^H]DA uptake, with a downward trend at 10 min and reductions to ∼50-60% of control values reached between 30-150 min post-injection (p<0.01-0.0001 vs. control) (**Figure 3A**). Previous characterization of this response demonstrated that transport activity remained suppressed for 3 h and returned to control levels by 24 h (Fleckenstein, Metzger et al. 1997, Kokoshka, Vaughan et al. 1998). Here we refined the recovery time, showing that activity remained suppressed between 4-6 h after injection and returned to control levels by 8 h (**Figure 3A**). Immunoblots of synaptosomes from these time points showed no changes in DAT protein levels (**Figure 3B**), indicating that transport reductions did not occur via loss of DAT protein. METH-induced reduction of transport thus persists for hours after palmitoylation has returned to control levels, supporting that events mediating resensitization of reuptake may not be strongly influenced by transporter palmitoylation status. The mechanism of transport reduction in these conditions was assessed via DA transport saturation analysis using synaptosomes prepared from striatal tissue collected 30 min after injection (**Figure 4**). Results show that transport losses were mediated by reductions in V_max_ from 47.4 ± 5.0 pmol/min/mg in synaptosomes from control animals to 38.0 ± 4.4 pmol/min/mg in synaptosomes from METH-treated animals (p<0.05), whereas there was no significant difference in K_m,DA_ between control (66.5 ± 3.6 nM) and METH conditions (63.7 ± 4.7 nM).

**Figure 4.**
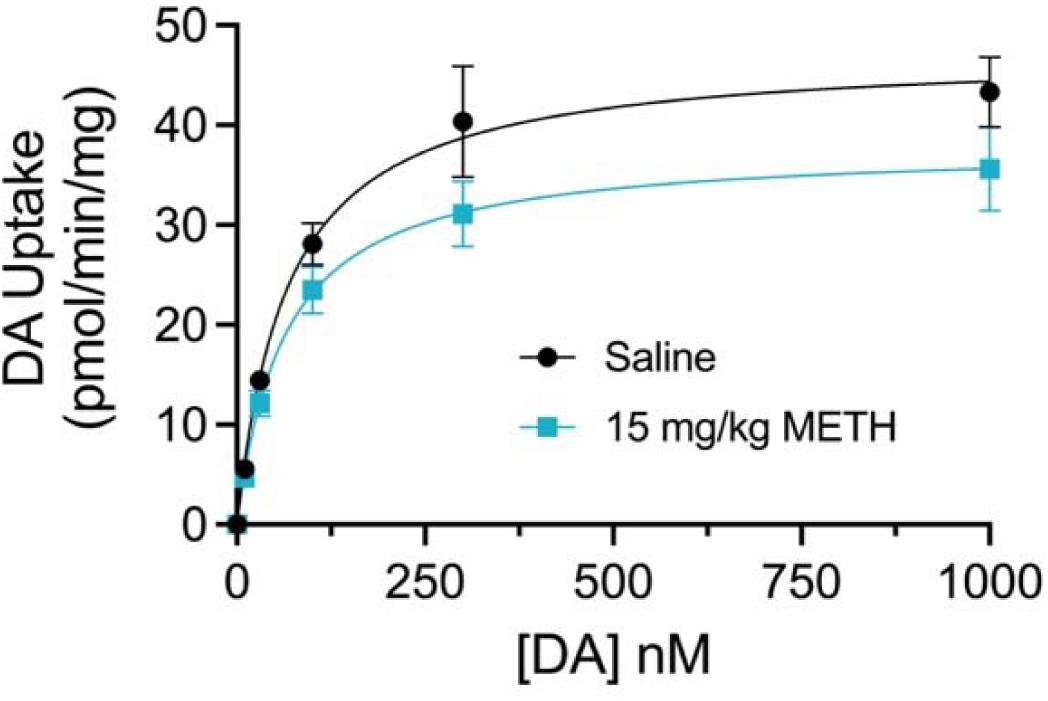
Kinetic analysis of DA transport activity after METH. Male Sprague-Dawley rats were given s.c. injections of saline or METH (15 mg/kg), sacrificed at 30 min, and synaptosomes subjected to DA transport saturation analysis. Curves show uptake activity (means ± S.E. from 4 independent experiments performed in triplicate). V_max_ and K_m_ values were determined in each experiment by nonlinear regression analysis using Michaelis–Menten kinetics with GraphPad Prism software. Mean V_max_ values were: saline 47.4 ± 5.0 pmol/min/mg; METH 38.0 ± 4.4 pmol/min/mg; mean K_m_ values were: saline 66.5 ± 3.6; METH 63.7 ± 4.7 nM.

We then examined the effects of *in vitro* METH on transporter palmitoylation using rDAT-LLCPK_1_ cells, which have been previously used to characterize DAT transport and phosphorylation responses to METH and PKC (Cervinski, Foster et al. 2005, Gorentla and Vaughan 2005) (**Figure 5**). In cells treated with 10 μM METH for 30 or 60 min, DAT palmitoylation was reduced to 83.2 ± 8.2% and 82.1 ± 3.8% of control, respectively (both p<0.05) (**Figure 5A**), whereas no reductions were seen in cells treated with 10 μM (−)-cocaine for 30 min (95.0 ± 5.2% of control, p>0.05) compared to reductions to 84.2 ± 4.1% of control by METH assessed in parallel (p<0.05) (**Figure 5B**). Although the reductions in palmitoylation induced by *in vitro* METH were generally of lesser magnitude than found *in vivo,* the responses are qualitatively similar and support that impacts on palmitoylation can occur via direct actions on DAT that do not require neuronal circuitry or transmitter signaling and are not induced by direct uptake blockade. Palmitoylation of DAT in this cell line is also reduced to a similar level by *in vitro* AMPH (not shown), indicating this effect as common to other psychostimulant substrates.

**Figure 5.**
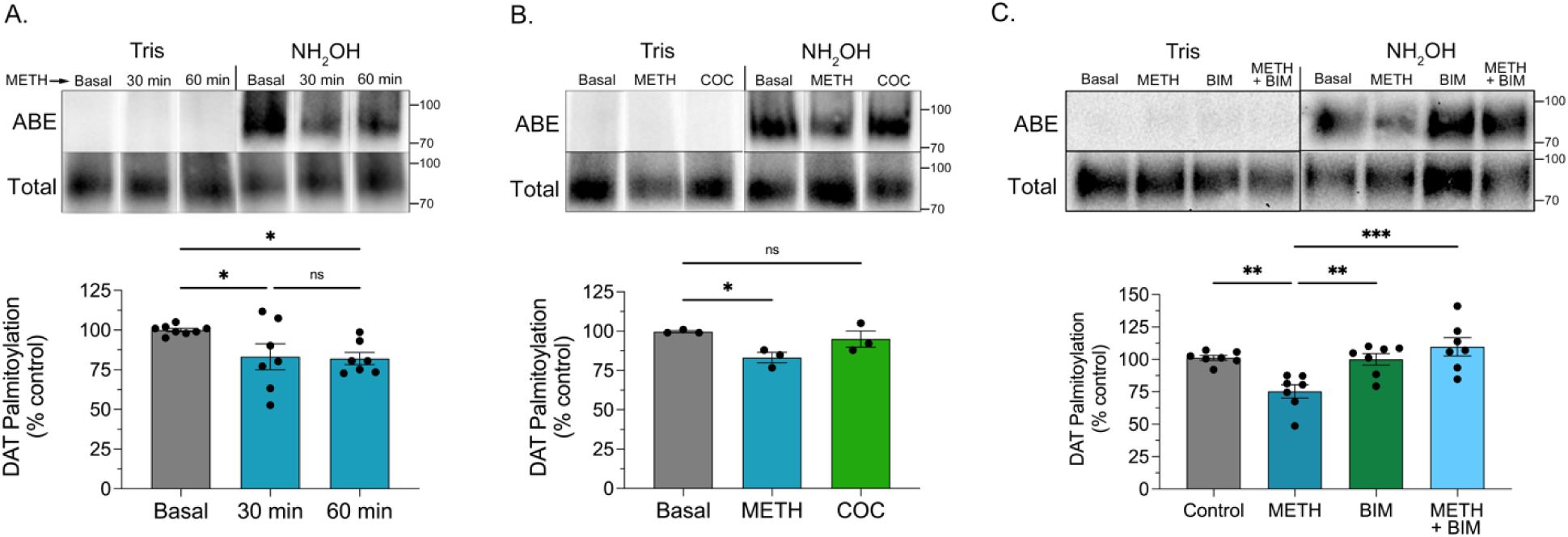
DAT palmitoylation is reduced by *in vitro* METH in a PKC-dependent manner. rDAT-LLCPK_1_ cells were treated with vehicle or METH (10 μM) for the indicated times (A), with (−)-cocaine (10 μM) or METH (10 μM) for 30 min (B), or with the indicated combinations of METH (10 μM) or BIM (10 μM) for 30 min (C), and membranes analyzed for DAT palmitoylation. Blots show representative ABE and Tris specificity controls (upper rows) and total DAT analyses (lower rows), and histograms show quantification of DAT palmitoylation normalized for total DAT and expressed as a fraction of vehicle control set to 100% (means ± S.E. of 3-7 independent experiments performed in duplicate). *, p<0.05; **, p<0.01, ***p<0.001 vs. indicated treatment groups, one-way ANOVA with Tukey’s post hoc test. n.s., not significant. White spaces between lanes in Panels A and B indicate removal of duplicate samples from the same experiment.

To test for involvement of PKC in these outcomes we pre-treated cells with the PKC inhibitor bisindolylmaleimide I (BIM) for 5 min followed by addition of METH and continued incubation for 30 min (**Figure 5C**). In these experiments, METH reduced DAT palmitoylation to 74 ± 5.1% of vehicle control (p<0.01), 10 μM BIM produced no effect (99.3 ± 4.5% of control, p>0.05), and addition of 10 μM BIM prior to and during METH treatment prevented reduction of palmitoylation (108.4 ± 6.9% of control, p>0.05). This supports the involvement of PKC in METH suppression of palmitoylation, which is mechanistically and functionally consistent with our earlier findings that palmitoylation is suppressed by PKC activation (Moritz, Rastedt et al. 2015) and that METH-induced transport down-regulation is PKC-dependent (Cervinski, Foster et al. 2005).

To determine the reversibility of DAT palmitoylation and down-regulation to *in vitro* METH we performed wash-out studies (**Figure 6**). Here rDAT-LLCPK_1_ cells were treated for 30 minutes with vehicle (gray bars) or 10 μM METH (blue bars), rapidly washed two times to remove drug, and either assayed immediately or incubated with buffer for 7 or 15 min prior to assessment. In palmitoylation studies (**Figure 6A**), modification of DAT in vehicle-treated cells was not changed in response to washing and incubations (all values p>0.05 vs. 0 min control). In METH-treated cells assayed immediately after washing, palmitoylation was reduced to 86.8 ± 7.5% of treatment-matched control, (p<0.05), similar to findings in Figure 5. With buffer incubation after removal of drug, palmitoylation recovered rapidly, returning to control levels within 7 min (96.0 ± 12.2% of control) and 15 min (96.5 ± 15.1%) (both p>0.05 vs. treatment-matched control). Whether this more rapid recovery of palmitoylation than seen in striatum follows from manual removal of drug compared to potentially slower metabolic clearance or represents a mechanistic difference between striatal and cell systems is not known.

**Figure 6.**
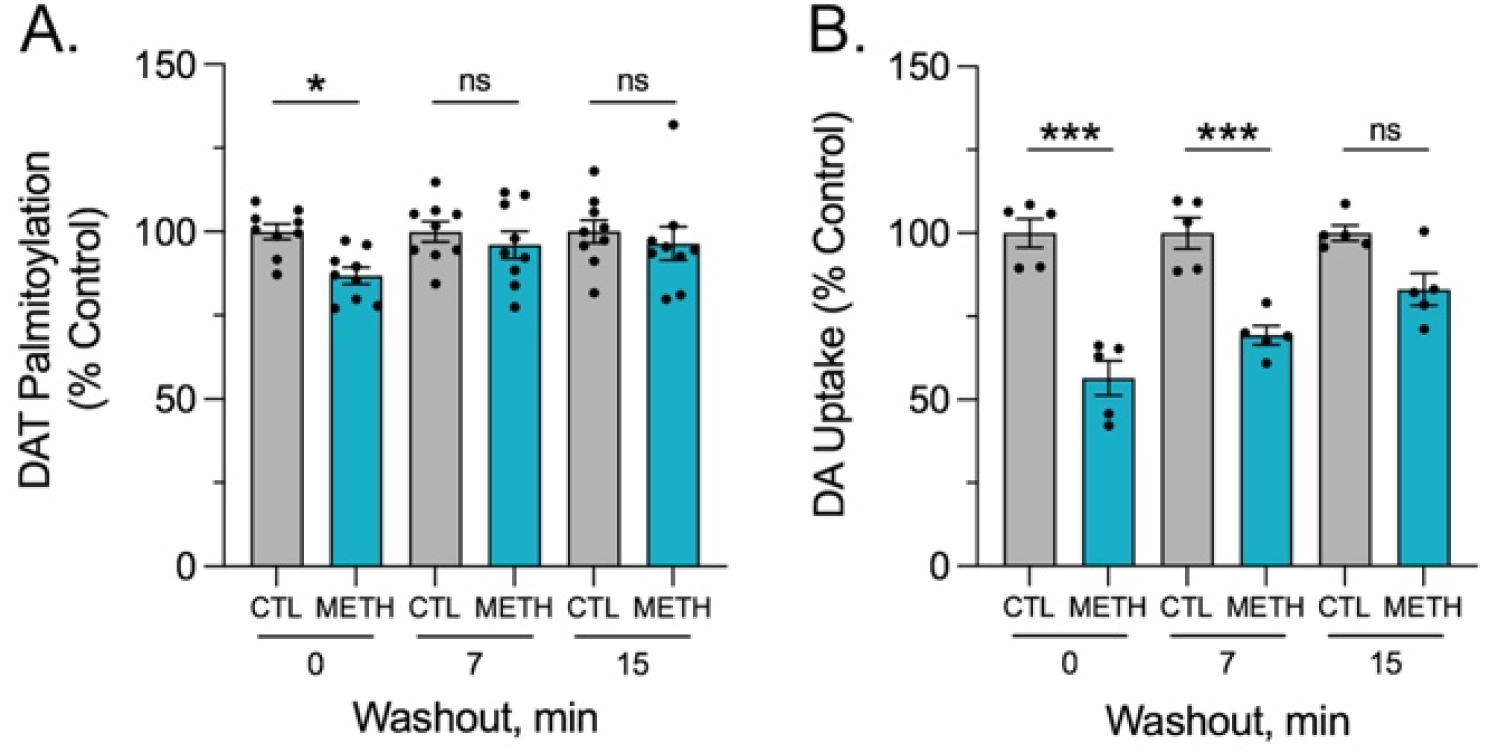
Reversibility of DAT palmitoylation and transport down-regulation to *in vitro* METH. rDAT-LLCPK_1_ cells were treated with vehicle (gray bars) or 10 μM METH (blue bars) for 30 min followed by washing to remove METH and incubation with buffer for the indicated times prior to analysis of palmitoylation or [^3^H]DA transport. (A). DAT palmitoylation normalized for total DAT protein at each time point and expressed as a fraction of treatment-matched controls set to 100% (means ± SE of 9 independent experiments). *, p<0.05 vs. control; one-way ANOVA with Fisher’s LSD test. (B). Transport activity expressed as a fraction of treatment-matched controls set to 100% (means ± S.E. of 5 experiments performed in triplicate). ***, p<0.001; one-way ANOVA with Tukey’s post hoc test. n.s., not significant.

Reversibility of *in vitro* METH effects on DA uptake is shown in **Figure 6B**. In cells that received vehicle treatment and washing (gray bars) transport activity showed no changes in magnitude at any time points (all values p>0.05 vs 0 min control). In METH-treated cells assessed immediately after washing, transport was reduced to 63.1 ± 10.4% of treatment-matched control (p<0.001), similar to down-regulation results seen in previous *in vitro* AMPH and METH pretreatment studies (Saunders, Ferrer et al. 2000, Cervinski, Foster et al. 2005). In treated and washed cells given incubations prior to analysis, transport remained down-regulated at 7 min (69.5 ± 5.6% of control, p<0.001), but recovered to 82.8 ± 7.9% of control level by 15 min (p>0.05) and to 100.9 ± 21.1% of control by 30 min (p>0.05) (not shown). This more rapid recovery of transport activity compared to that seen *in vivo* again potentially reflects manual versus metabolic clearance of drug or mechanistic differences between systems, but qualitatively parallels findings from striatum that transport can remain down-regulated after palmitoylation has returned to starting levels.

About 50% of endogenous palmitate incorporation on rDAT occurs on residue Cys580 at the base of TM12, with modification of this site mechanistically linked to increased transport velocity and suppression of PKC-induced down-regulation (Foster and Vaughan 2011). To further investigate palmitoylation contributions to METH-induced down-regulation we directly compared down-regulation responses of C580A and WT DAT to METH treatment (**Figure 7A**). Here WT- and C580A-rDAT cells were treated in parallel with 10 μM METH for the indicated times, washed to remove the drug, and assayed for uptake. Similar to previous findings (Cervinski, Foster et al. 2005), transport activity of WT DAT displayed significant down-regulation within 2 min of drug application, with activity plateauing at ∼70% of starting levels (all values p<0.001 vs. WT DAT control). In assays performed in parallel, transport activity of C580A DAT showed similar rapid reductions, with activity plateauing at ∼50% of the C580A starting value (all values p<0.001 vs. C580A DAT control). At all time points, the magnitude of C580A DAT down-regulation was significantly greater than that of WT DAT (all values p<0.05-0.01 vs. time-matched WT DAT), supporting that METH-induced transport down-regulation is opposed by palmitoylation of Cys580.

**Figure 7.**
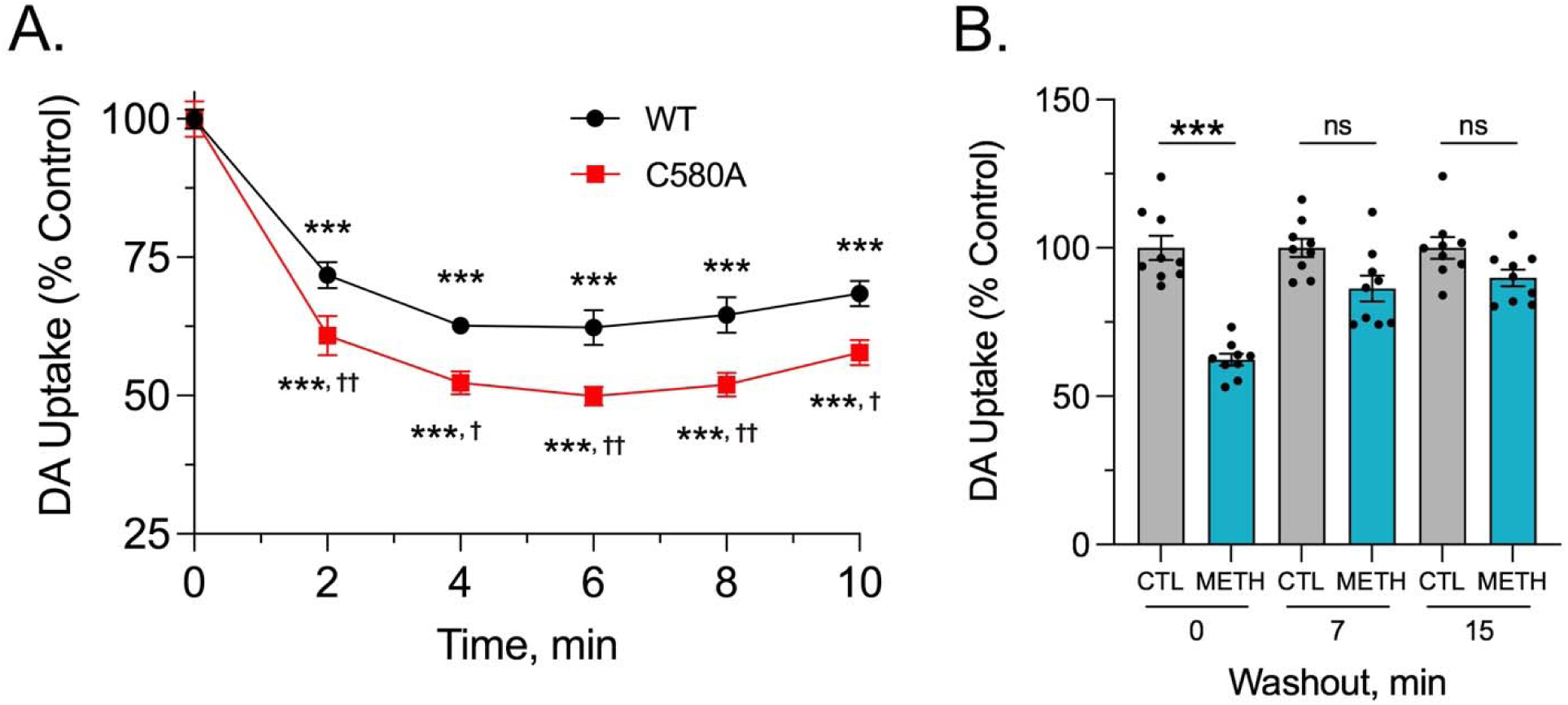
C580A DAT undergoes enhanced down-regulation to METH and rapid reversibility after METH washout. (A). LLPCK_1_ cells expressing WT rDAT (black) or C580A rDAT (red) were treated with vehicle or 10 μM METH for the indicated times, washed to remove METH, and assayed for [^3^H]DA uptake. Symbols represent means ± S.E. of 3 experiments performed in triplicate. ***, p<0.001 METH vs. respective vehicle controls for each form. ^†^, p<0.05; ^††^, p<0.01; C580A DAT vs. WT DAT at matching time points, one-way ANOVA with Fisher LSD test. (B). C580A-rDAT-LLCPK_1_ cells were treated with vehicle (gray bars) or 10 μM METH (blue bars) for 30 min followed by rapid washing and incubation with buffer for the indicated times prior to analysis of [^3^H]DA transport. Histogram shows transport activity of control and treated cells at each time point after washing (means ± S.E. of 9 experiments performed in triplicate), expressed as a fraction of vehicle control for each time point set to 100%. *** p<0.001; one-way ANOVA with Tukey’s post hoc test. n.s., not significant.

The reversibility of METH-induced C580A DAT down-regulation was examined using the same washout paradigm as for WT DAT, with C580A DAT cells treated with vehicle or 10 μM METH for 30 min followed by washout and incubation prior to analysis of uptake activity (**Figure 7B**). Transport activity in cells given vehicle treatment showed no changes after washing and incubation (all p>0.05 vs 0 min control). In METH-treated cells that were assayed immediately after washing, transport of C580A DAT showed reductions to 62.3 ± 6.0% of control (p<0.001), similar to that shown in Fig. 7A. In cells given buffer incubations after washing, uptake activity returned to control levels by 7 min (86.3 ± 13.1% of control) and 15 min (89.9 ± 8.4% of control) (both p>0.05 vs time-matched control), indicating that C580A DAT transport activity recovered rapidly after removal of METH. Whether this rate of recovery differs from that of WT DAT was not directly investigated.

## DISCUSSION

The pathophysiological dysregulation of DAT by METH implicates the underlying processes in neurochemical outcomes and as potential targets for therapeutic intervention in drug addiction and other reuptake disorders. The findings shown here now extend our understanding of METH mechanisms by demonstrating a previously unknown effect on transporter palmitoylation that shares time course, PKC dependency, and circuit/receptor-independence characteristics with METH-induced transport down-regulation. Because the velocity of DAT is enhanced by palmitoylation, these findings, in conjunction with the elevated down-regulation of the palmitoylation-deficient form, support that the modification serves to oppose the acute down-regulation of transport induced by METH that is associated *in vivo* with hyperdopaminergia.

Mechanistically the suppression of down-regulation by palmitoylation can be envisioned as the modification serving to stabilize higher velocity DAT forms or complexes such that reduced modification enables or enhances the transition of transporters into lower velocity states, as shown schematically in **Figure 8**. The precise mechanisms by which this might occur are not known, but down-regulation and endocytosis of DAT induced by AMPH, METH, and PKC involve many similar processes that could be impacted by palmitoylation, as described below. Several studies support that regulation of DAT occurs by simultaneous kinetic and trafficking processes that may display different onset and reversal time courses, and in this study we have not attempted to distinguish between these regulatory modes. Specific events may also differ or involve additional regulatory components, depending on stimulation conditions or systems analyzed (Torres 2006, Eriksen, Jorgensen et al. 2010, Schmitt and Reith 2010, Bermingham and Blakely 2016, Sorkina, Cheng et al. 2021, Bolden, Pavchinskiy et al. 2025), which might also relate to these issues.

**Figure 8.**
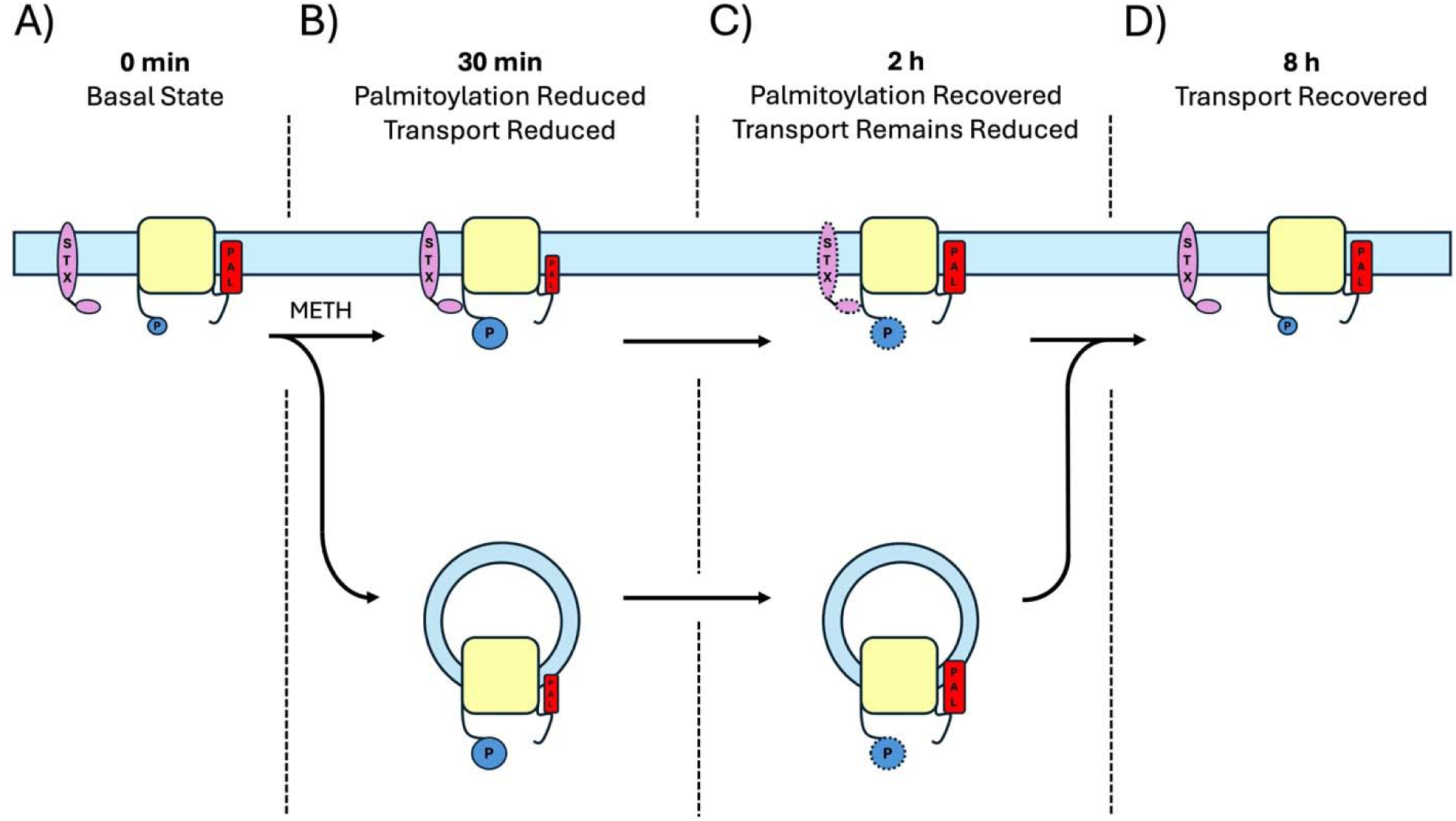
Model depicting potential events in METH-induced responses of DAT palmitoylation and down-regulation in rat striatum. DAT populations are shown in four functional and biochemical states prior to and after injection of METH. A). In the basal state DAT is shown at the plasma membrane (blue line) with high levels of palmitoylation (red rectangles) and low levels of phosphorylation (blue circles) that synergistically contribute to baseline reuptake capacity. B). 30 min after METH injection DAT palmitoylation is reduced and transport is down-regulated. Potential mechanisms leading to reduced reuptake are increased transporter phosphorylation, increased interaction with STX (purple oval), and/or removal of DAT from the surface via endocytosis (vesicles). For simplicity, other structural, interactome, or membrane mechanisms that could mediate down-regulation as described in the text are not shown. C). 2 h after METH injection, DAT palmitoylation has returned to initial levels but transport activity remains reduced. Events that could potentially underlie this condition are retention of high phosphorylation and/or-STX interactions (indicated by stippling), or retention of endocytosed DAT in cytoplasmic vesicles. D). 8 h after METH injection, DA transport has returned to initial levels. Events that could potentially underlie this response are dephosphorylation of transporters to initial levels, dissociation of DAT-STX interactions, and return of endocytosed transporters to the plasma membrane.

The regulatory property with the most current evidence for interfacing with both METH and palmitoylation is N-terminal phosphorylation, which kinetically suppresses reuptake velocity of surface transporters (Moritz, Rastedt et al. 2015), is stimulated within minutes of *in vivo* or *in vitro* METH (Cervinski, Foster et al. 2005, Moritz, Rastedt et al. 2015, Challasivakanaka, Zhen et al. 2017), and is inhibited by palmitoylation (Moritz, Rastedt et al. 2015). Suppression of phosphorylation by palmitoylation can occur in the absence of exogenous kinase or phosphatase modulation, supporting a direct connection between reduced palmitoylation and higher phosphorylation/lower velocity states.

A potentially congruent mechanism is binding of the SNARE protein syntaxin 1A (STX) to DAT, which occurs through the transporter N-terminal domain (Lee, Kim et al. 2004, Binda, Dipace et al. 2008). In addition to regulating DAT channel and efflux properties (Carvelli, Blakely et al. 2008, Shekar, Mabry et al. 2023), STX suppresses DA transport velocity and stabilizes the tonic level of transporter phosphorylation (Cervinski, Foster et al. 2010). Binding of STX to DAT is increased by AMPH (Binda, Dipace et al. 2008), consistent with drug-induced stimulation of transporter phosphorylation and down-regulation. The impact of DAT palmitoylation on STX mechanisms has not yet been investigated, but STX is also palmitoylated (Vardar, Salazar-Lazaro et al. 2022), which modulates its membrane fusion properties and could potentially influence its DAT interactions or outcomes.

Palmitoylation also impacts multiple DAT membrane properties that have been associated with transport regulation and endocytosis. Computational modeling studies support that palmitoylation of hDAT Cys581 promotes structural conformations that enhance homodimerization (Zeppelin, Pedersen et al. 2021), which has been linked to reuptake activity and membrane trafficking (Cheng, Garcia-Olivares et al. 2017, Zhen and Reith 2018, Sorkina, Cheng et al. 2021). In addition, DAT lateral membrane mobility, which may influence microdomain targeting or regulome interactions (Sorkina, Doolen et al. 2003, Zeppelin, Ladefoged et al. 2018, Shekar, Mabry et al. 2023), is reduced by palmitoylation and increased by phosphorylation, PKC, and AMPH (Shetty, Bolland et al. 2023). For many proteins palmitoylation also influences regulatory events related to membrane cholesterol (Lorent and Levental 2015), and DAT kinetic properties, phosphorylation, and endocytosis have all been linked to cholesterol and transporter partitioning into cholesterol-rich membrane raft domains (Foster, Adkins et al. 2008, Hong and Amara 2010, Cremona, Matthies et al. 2011, Jones, Zhen et al. 2012), although a direct role for palmitoylation in these events has not yet been directly examined.

Possible impacts of palmitoylation on the transporter C-terminal domain are as yet speculative, but the location of Cys580/581 suggests the potential for the modification to alter TM12 properties such as conformation, tilt, membrane hydrophobic matching, or hydrophobic protein-protein interactions. Such impacts could be propagated to the C-terminus, including the adjacent juxtamembrane helix between residues 587-592 that is proposed to modulate transporter kinetics via interaction with intracellular loops (Penmatsa, Wang et al. 2013). This helix is also necessary for PKC-stimulated endocytosis and binds the trafficking-regulatory proteins G_βγ_ and Rit2 (Holton, Loder et al. 2005, Navaroli, Stevens et al. 2011, Fagan, Kearney et al. 2020, Pino, Nuñez-Vivanco et al. 2021), with a Rit2 homolog mediating AMPH behavioral sensitivity in *Drosophila* (Fagan, Kearney et al. 2021).

Less is known about the events that occur after METH clearance that underlie DAT resensitization and return of the reuptake system to homeostasis. The longer time needed for recovery of uptake compared to palmitoylation indicates that METH down-regulation mechanisms reverse more slowly and may remain engaged with repalmitoylated transporters. This could occur via persistence of phosphorylation or STX binding with repalmitoylated surface transporters, or by endocytosed transporters undergoing repalmitoylation prior to plasma membrane recycling, as shown schematically in **Figure 8**, or by sustained involvement of repalmitoylated DATs with one or more of the other potential regulatory conditions described above.

How METH exposure results in reduction of DAT palmitoylation is not known. Protein palmitoylation is catalyzed by palmitoyl acyltransferases (PATs) and depalmitoylation is catalyzed by acyl protein thioesterases (APTs) and protein palmitoyl thioesterases (PPTs) (Resh 2006, Jin, Zhi et al. 2021), such that reduction of DAT palmitoylation could follow from reduced activity of PATs and/or enhanced activity of APTs or PPTs (**Figure 9**). These enzymes are themselves under posttranslational control (Jin, Zhi et al. 2021), and acute dysregulation of their actions could potentially follow from changes in PKC or other regulatory enzymes through METH-induced alterations in cytoplasmic Ca^2+^ (Giambalvo 1992, Gnegy, Khoshbouei et al. 2004). To date five PATs that enhance DAT palmitoylation have been identified (Bolland, Moritz et al. 2019), suggesting these proteins as potential loci for involvement in these events. In addition to events driven by changes in enzyme activity, a non-exclusive possibility is that drug-induced acceleration of transport could alter palmitoylation levels by impacting modification site accessibility or regulome constituents.

**Figure 9.**
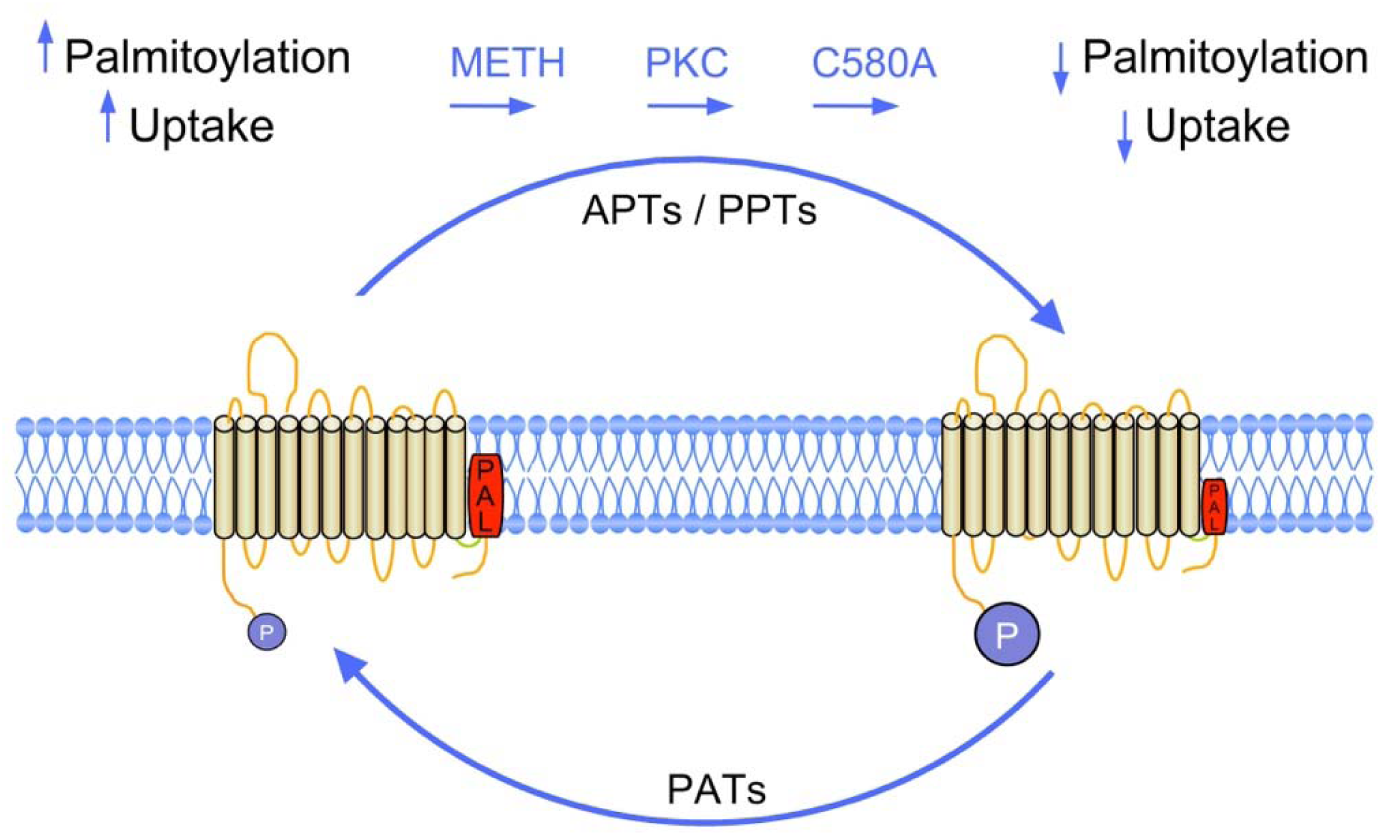
Model of DAT regulation by palmitoylation and METH. DAT exists in populations possessing lower or higher levels of palmitoylation (red rectangles) driven by the actions of PAT and APT/PPT enzymes, with phosphorylation (blue circles) undergoing reciprocal responses as indicated. In the absence of exogenous stimulation (left) the tonic level of DAT palmitoylation is high and phosphorylation is low, promoting a form that possesses higher transport V_max_. In the presence of METH, activation of PKC, or C580A mutation, palmitoylation is reduced, phosphorylation is increased, and transport V_max_ is reduced, (right).

This study has described a dysregulatory DAT palmitoylation response following an acute exogenous stimulus, but the findings support the potential for involvement of transporter palmitoylation in additional conditions such as long-term drug use, endogenous reuptake disorders, or regional/sex-specific transporter regulation. DAT displays pleiotropic responses to palmitoylation, with acute alterations linked to rapid transport regulation as described in this study, whereas chronic suppression leads to enhancement of transporter degradation and steady-state reductions in total levels (Bolland, Moritz et al. 2019). As such, the transient suppression of DAT palmitoylation by acute METH suggests the potential for chronic METH to induce long-term reductions in palmitoylation that lead to the reduced levels of transporter expression and reuptake capacity that are hallmarks of METH abuse (Volkow, Chang et al. 2001, Jayanthi, Daiwile et al. 2021).

Palmitoylation could potentially also be impacted by endogenous conditions that contribute to DA imbalance disorders or establish differential regulatory baselines. Many PAT, APT, and PPT enzymes are subject to mutations and expression deficits that result in improper modification and function of target substrates. In neuronal systems these substrates include major synaptic constituents such as receptors, channels, SNAREs, and scaffolds, with dysregulated palmitoylation associated with diseases including schizophrenia, attention deficit hyperactivity disorder, and major depressive disorder (Liao, Huang et al. 2023, Peng, Liang et al. 2024), supporting that similar conditions could contribute to dysregulated DAT modification and function in DA disorders. Dysregulation of DAT palmitoylation could also represent a mechanism underlying the anomalous kinetics that have been observed in transporter polymorphic variants associated with DA imbalance disorders. In this regard, we have found that hDAT isoform A559V identified from individuals diagnosed with attention deficit hyperactivity disorder and bipolar disorder (Mazei-Robison, Bowton et al. 2008, Bowton, Saunders et al. 2014) displays tonically reduced *in vitro* palmitoylation (unpublished result) that could relate to its altered kinetic properties, AMPH insensitivity, and enhanced phosphorylation. Increasing evidence also supports that tonic regulation of DAT can differ between brain regions and sexes that result in differential neurochemical and behavioral outcomes (Gowrishankar, Gresch et al. 2018, Stewart, Mayer et al. 2022, Mayer, Stewart et al. 2024). In some cases these differences have been linked to dysregulated transporter phosphorylation, which, by extension, supports potential involvement of palmitoylation.

The modification of DAT by palmitoylation thus represents a key component of the transporter regulatory machinery that could function in spatial and temporal control of DA clearance in normal and pathophysiological states. Issues for further elucidation to inform understanding of pathogenesis and potential therapeutic interventions include determining the mechanisms of palmitoylation in kinetic functions, membrane trafficking, and responses to genetic conditions.

## Acknowledgments

This work was supported by the University of North Dakota and grants ND EPSCoR Doctoral Dissertation Fellowship OIA-1355466 (MJH); P20 GM104360 (to UND) from the COBRE program and P20 GM103442 (to UND) from the INBRE program of the National Institute of General Medical Sciences. LC-MS analysis was performed by the MS Core Facility at the University of North Dakota supported by the School of Medicine and Health Sciences.

## Author Contributions

RAV and JDF conceived experiments, analyzed data, and wrote the paper. MJH, DEB, CDK, MS, ACB, MYG, SAG, and CRB performed experiments, analyzed data, and assisted in manuscript writing.

## Data Availability

The data that support the findings of this study are included in the Materials, Methods, and Results sections of this article.

## Assurances

All procedures involving animals were approved by the University of North Dakota Institutional Animal Care and Use Committee in accordance with the National Institutes of Health Guide for the Care and Use of Laboratory Animal Animals. Experiments involving radioactivity were approved by the University of North Dakota Radiation Safety Committee, and experiments involving recombinant transporters were approved by the University of North Dakota Institutional Biosafety Committee.

## Abbreviations

ABE: acyl biotinyl exchange
AMEM: Alpha minimal essential medium
AMPH: amphetamine
APT: acyl protein thioesterase
BCA: bicinchoninic acid
BIM 1: bisindolylmaleimide 1
DA: dopamine
EDTA: ethylenediaminetetraacetic acid
HEPES: 4-(2-hydroxyethyl)-1-piperazineethanesulfonic acid
HPDP biotin: sulfhydryl-reactive (*N*-(6-(biotinamido)hexyl)-3′-(2′-pyridyldithio)-propionamide
KRH: Krebs-Ringers-HEPES buffer
LLC-PK_1_: Lily Lung Carcinoma Porcine Kidney Cells
METH: methamphetamine
NA: hydroxylamine
NEM: N-ethylmaleimide
PAT: palmitoyl acyltransferase
PKC: protein kinase C
PMSF: phenylmethanesulfonyl fluoride
PPT: palmitoyl protein thioesterase
RIPA: radioimmunoprecipitation assay
SP: sucrose phosphate buffer

